# Dissecting Autism Genetic Risk Using Single-cell RNA-seq Data

**DOI:** 10.1101/2020.06.15.153031

**Authors:** Siying Chen, Xueya Zhou, Eve Byington, Samuel L. Bruce, Haicang Zhang, Yufeng Shen

## Abstract

Autism spectrum disorder (autism) is a condition with strong but heterogenous genetic contribution. Recent exome and genome sequencing studies have uncovered many new risk genes through *de novo* variants. However, a large fraction of enrichment of *de novo* variants observed in cases are not accounted for by known or candidate risk genes, suggesting that the majority of risk genes are still unknown. Here we hypothesize that autism risk genes share a few common cell-type specific gene expression patterns during brain development, and such information can be quantified to improve statistical power of detecting new risk genes. We obtained large-scale single-cell RNA-seq data from human fetal brain collected through a range of developmental stages, and developed a supervised machine-learning approach “A-risk” (Autism risk), to predict the plausibility of autism risk genes across the genome. Using data from recent exome sequencing studies of autism, A-risk achieves better performance in prioritizing *de novo* variants than other methods, especially for genes that are less intolerant of loss of function variants. We stratified genes based on A-risk and mutation intolerance metrics to improve estimation of priors in extTADA and identified 71 candidate risk genes. In particular, *CLCN4, PRKAR1B, and NR2F1* are potentially new risk genes with further support from neurodevelopmental disorders. Expression patterns of both known and candidate risk genes reveals the important role of deep-layer excitatory neurons from adult human cortex in autism etiology. With the unprecedented revolution of single-cell transcriptomics and expanding autism cohorts with exome or genome sequencing, our method will facilitate systematic discovery of novel risk genes and understanding of biological pathogenesis in autism.

## Introduction

Autism spectrum disorder (autism) is a phenotypically heterogeneous developmental disorder, affecting 1 in 59 children in the United States [1]. Earlier studies have shown a strong genetic basis for autism with up to 90% concordance between monozygotic twins [2, 3] and 10-fold higher chance for younger sibling to be diagnosed with autism if there is an older affected sibling [4, 5]. Simulations estimate one thousand autism risk genes with large effect [6]; however, currently only about 100 known risk genes [7] have robust evidence from recent studies [6, 8, 9]. These known risk genes only account for less than 5% of autism cases [10]. Therefore, it is critically important to identify new risk genes. However, the identification of new risk genes based on statistical evidence is limited by lack of power due to sample sizes.

A general approach to improve the power for detecting risk genes is to use prior knowledge and functional genomic data to predict plausibility of candidate risk genes. Previous studies have implemented network-based methods utilizing genotype-phenotype associations [1, 11, 12], protein-protein physical interactions [13], brain-specific functional interactions [14] and gene coexpression networks [15, 16]. We previously developed a semi-supervised method using cell-type specific expression profiles from mouse bulk microarray data based on Principle Component Analysis (PCA) [17]. One advantage of using cell-type specific expression is the ability to jointly infer plausible risk genes and cell types that are correlated with risk plausibility, potentially improving the understanding of the disease mechanism. Our method was limited by the lack of spatiotemporal cell-type information from developing brains and the species difference between mouse and human. Recent studies have developed machine learning approaches to classify autism risk genes with human brain expression data [18, 19], but are still limited by the resolution of data in cell types or developmental stages pertinent to the disease.

With the motivation to identify new risk genes for autism, here we developed a supervised machine learning method based on gradient boosting trees, “A-risk” (Autism risk), that can learn known risk genes’ expression patterns in single-cell transcriptomics of human fetal midbrain and prefrontal cortex, to then predict the plausibility of any gene being an autism risk gene. We hypothesize that autism risk genes have distinct spatiotemporal expression signatures in developing human brain in neurotypicals. When comparing A-risk to other metrices or methods in prioritizing risk variants, we observed better performance of A-risk in prioritizing candidate risk variants using *de novo* variant data of 8838 trios from recent publications[6, 20–24]. Furthermore, we showed that A-risk and gene mutation intolerance metrics[25] can be combined to improve prior estimation in an empirical Bayesian model and enables identification of additional novel risk genes. Finally, we investigated the cell type specific expression patterns in adult brain of known and novel autism risk genes and found that they are highly expressed in deep-layer excitatory neurons in adult human cortex, suggesting the association of deep excitatory neurons in cortex to the etiology of autism.

## Results

### Single-cell expression pattern is correlated with autism risk

We obtained two single-cell RNA-seq data sets from human fetal midbrain and prefrontal cortex. The midbrain data are mostly from the first trimester [26], while the prefrontal cortex data are mostly from the second trimester [27]. Previous studies have suggested the role of prefrontal cortex [28–31] and midbrain dopamine system [32–34]. On average, 2302 and 4503 genes per cell are detected in the midbrain and the prefrontal cortex data, respectively (Supplementary Figure 1). We obtained the cell type labels from original publications, and then define the expression level of a gene in a cell type as the fraction of cells with ≥1 UMIs (Unique Molecular Identifiers) in the cell type at a certain developmental time point. The feature set of our data is the combination of cell types and developmental time points (Supplementary Table 1).

To investigate temporal and cell type specific expression pattern of autism risk genes, we collected 88 known autism risk genes from the SFARI (Simons Foundation Autism Research Initiative) Gene database [7] (released version on 08/29/2019, score 1 or 2), which are genes strongly implicated in autism based on expert curation from the literature. We also obtained 154 genes with at least 1 *de novo* LGD (likely-gene disrupting) variant in unaffected siblings from an exome-sequencing study[6] (Supplementary Table 2), representing non-risk genes with random *de novo* mutations. Known risk genes tend to have a wide range of average expression level in both data sets, while non-risk genes have lower average expression (Supplementary Figure 2A). We performed PCA (Principle Component Analysis) of these two groups of genes using expression level from the single cell data sets. The first component partially separates known risk genes and non-risk genes (Supplementary Figure 2B). This is consistent with previous findings using bulk RNA microarray data from mouse brain [17].

To leverage the temporal and cell type specific expression pattern of known autism risk genes, we developed a supervised machine learning method, “A-risk”, to predict plausibility of being an autism risk gene for all protein-coding genes (Supplementary Table 3). A-risk is based on gradient boosting. We train the model using 88 known autism risk genes as positives and the 154 non-risk genes as negatives. Figure 1A shows the overall workflow of A-risk. Five-fold cross-validation during training achieves an average AUC (Area Under Curve) of ROC (Receiver Operating Characteristic) curves at 0.77 (Supplementary Figure 3A). A-risk score distribution shows a large separation of known risk genes and non-risk genes (Figure 1B). We chose A-risk 0.4, corresponding to top 2642 ranked genes, as a recommended cutoff for analysis where a binary stratification of genes is needed.

**Figure 1.**
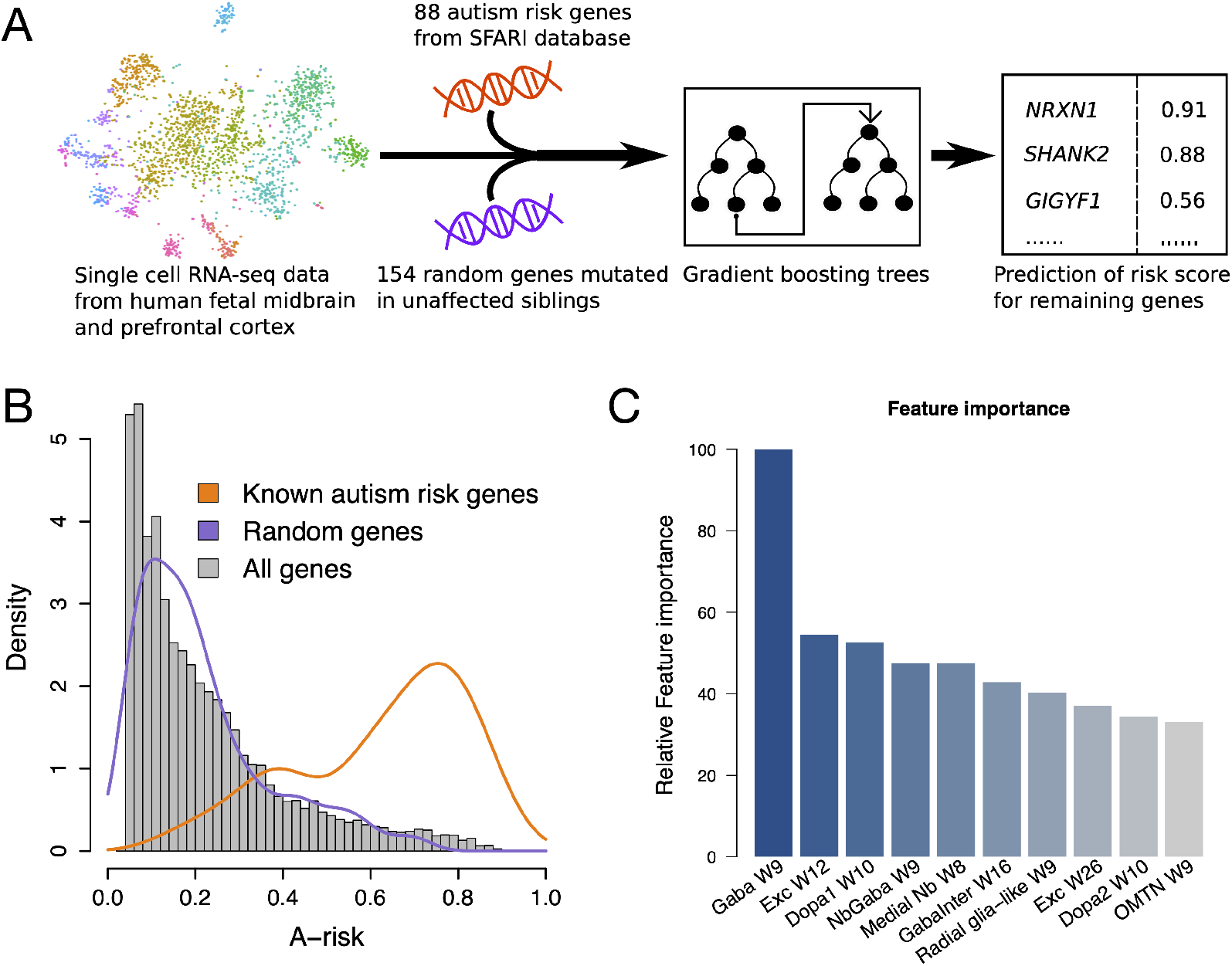
A-risk, a gradient boosting tree model to estimate plausibility of being risk genes of autism from single-cell RNA-seq data. (A) A flowchart of the method. (B) A-risk score distribution. A-risk of all genes in the genome are shown in the histogram in gray. The distribution of A-risk of known autism risk genes and randomly mutated genes, which are positive and negative training sets in A-risk model respectively, are shown as orange and purple density curves. A-risk score 0.4 is where the positives and negatives show separation. (C) “Feature importance” derived from the gradient boosting trees model showing cell types from both midbrain late first trimester and prefrontal cortex second trimester make substantial contribution to the prediction. The y-axis is the relative important of each feature against the max, which is GABAergic neurons in midbrain at week 9. W, week. Gaba, GABAergic neurons. Exc, excitatory neurons. Dopa, Dopaminergic neurons. NbGaba, neuroblast GABAergic. Nb, neuroblast. GabaInter, GABAergic interneurons. OMTN, oculomotor and trochlear nucleus.

We quantify the contribution of cell types to A-risk prediction by feature importance, a score for each feature measuring how valuable it is in constructing the model. The top ranked cell types are GABAergic neurons in midbrain at week 9, dopaminergic neurons in midbrain at week 10 and prefrontal cortex excitatory neurons at week 12 (Figure 1C). Overall, cell types from both midbrain late first trimester and prefrontal cortex second trimester made substantial contribution to the prediction. The full list of feature importance from the model is available in Supplementary Table 4.

### A-risk improves prioritization of *de novo* variants in autism cases

To investigate if A-risk can prioritize *de novo* risk variants detected from exome or genome sequencing studies, we compiled *de novo* likely gene-disrupting (LGD) variants of 8838 trios from recent published studies [6, 20–24] (Supplementary Table 5). We calculated enrichment rate of LGD *de novo* variants in a gene set by the observed number of variants divided by the expected number estimated from background mutation rate models [35, 36] (Table 1). The enrichment rate for all genes excluding known risk genes is 1.4, suggesting there are additional risk genes that harbor *de novo* LGD variants. When further selecting genes by A-risk ≥0.4, the enrichment rate reaches 2.1 (p-value=1.3e-32, Poisson test), showing that A-risk can increase the signal-to-noise ratio in prioritized candidate risk genes.

**Table 1.** A-risk improves prioritization of *de novo* LGD variant in autism cases (n=8836).

|  | Observed number of variants | Expected | Enrichment Rate | P-value |
| --- | --- | --- | --- | --- |
| All genes (N=18663) | 1341 | 784 | 1.7 | 3e-73 |
| Excluding known risk genes (N=18575) | 1114 | 774 | 1.4 | 9e-31 |
| A-risk$\geq 0.4$, excluding known risk genes (N=2566) | 313 | 148 | 2.1 | 1e-32 |

To further assess the utility of A-risk in prioritizing novel risk genes, we compute enrichment and precision-recall like curves and compare with other methods. The precision-recall like curves compare the utility of each method in prioritizing true risk variants[35, 36]. With each method, we rank all genes. In all genes above a certain rank threshold, we estimate the number of detected true risk variants (“positives”) by the difference of observed number of variants (“detected positives”) and expected number. The total number of true positives is unknown, but it is a constant independent of methods. Therefore, the estimated number of true positives is a proxy of recall. The estimated precision is the number of detected true positives divided by the total number of detected positives. Besides the *de novo* LGD variants we used for Table 1, we included deleterious missense (D-mis) variants defined by REVEL score [37] ?0.5 in the following analysis. In addition, all known risk genes used in model training are excluded from analysis. We compared A-risk with mouse brain bulk expression ranks at E9.5 [38], ExAC pLI [25], and the baseline where the corresponding estimates are calculated in all protein-coding genes (excluding known risk genes). A-risk achieves consistently higher enrichment from the top 2000 to top 4000 ranked genes compared to others and significantly higher than the genome baseline (Figure 2A). At the 2500 top rank, roughly corresponding to A-risk score 0.4, A-risk achieves better precision than other metrices and prioritizes almost half of total *de novo* variants with a relatively high precision (0.46), a 64% improvement from the genome-wide baseline (precision=0.28) (Figure 2B). Furthermore, in non-constrained genes (pLI<0.9), A-risk shows significantly higher enrichment and better precision compared to mouse brain expression levels (Figure 2C and D), indicating A-risk is complementary to pLI with the potential to optimize risk gene discovery, especially among non-constraint genes. We also compared A-risk with other recent methods aimed to find novel autism risk genes, such as D-score [17] and Krishnan 2016 [14] (Supplementary Figure 4). A-risk again shows superior performance in enrichment, precision and true positives from top 1500 to top 4000 ranks of the three methods (Supplementary Figure 4A and B), and particularly in non-constrained genes (Supplementary Figure 4C and D).

**Figure 2.**
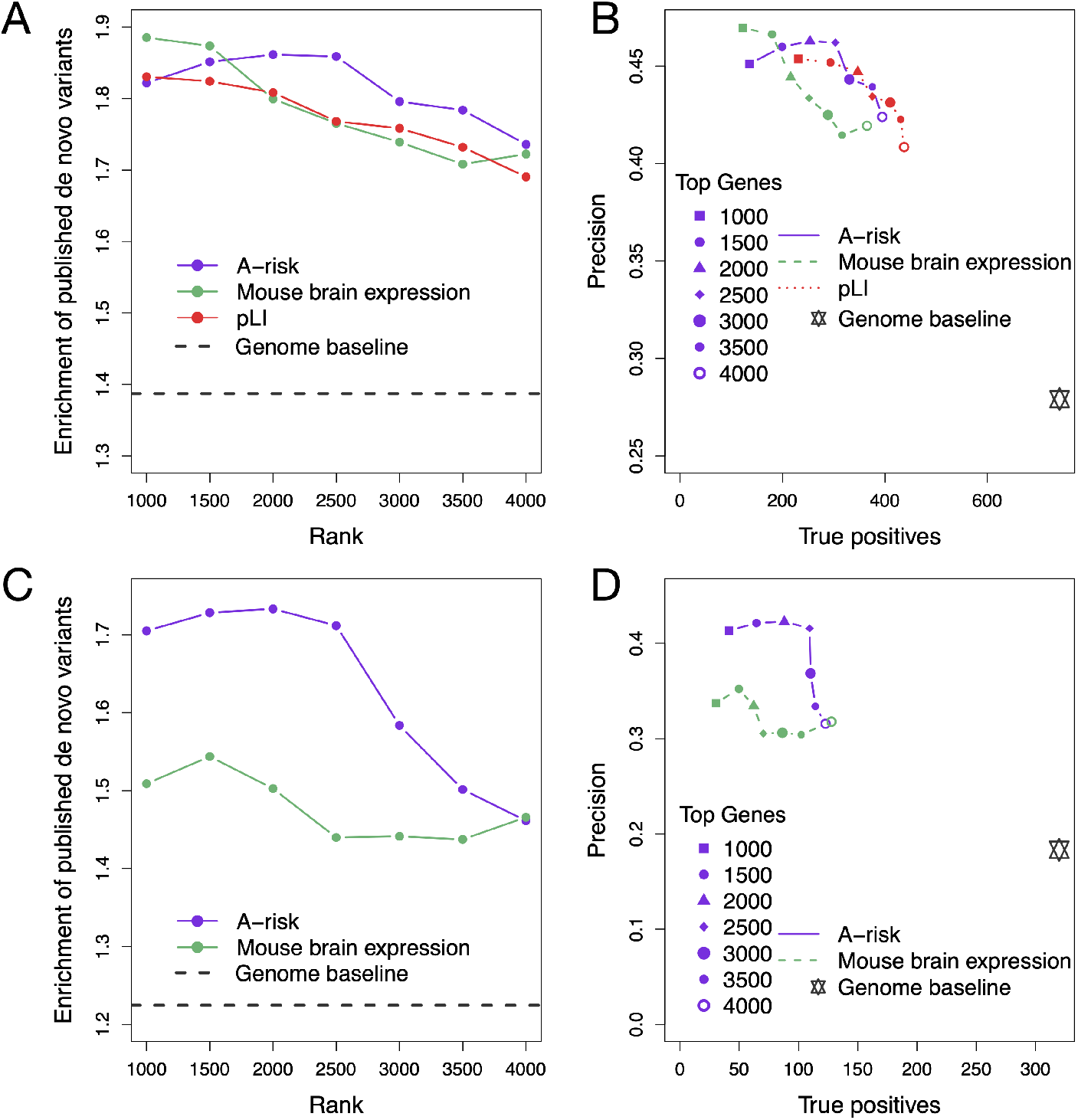
Superior performance of A-risk in prioritization of *de novo* variants at top 2500 ranks, especially in non-constraint genes. A-B, comparison of A-risk to mouse brain expression level, pLI and genome baseline in prioritization of *de novo* LGD and D-mis variants among top genes ranked by each individual metrics, excluding known risk genes used in A-risk training. D-mis is defined by REVEL score ? 0.5. The *de novo* variant data is compiled from 8838 published trios of exome sequencing studies. (A) Enrichment is the ratio of observed number of *de novo* variants to the expected number of *de novo* variants estimated by background mutation rate in top ranks, ranging from top 1000 to top 4000 genes. (B) Precision and true positives compared in top ranks. True positives, which are the difference value between observed number of de novo variants and the expected number, represent the recall since the true number of total causal variants is unknown. Precision is computed as dividing true positives by the observed number. Genome baseline is the grey star in the plot. C-D, comparison of A-risk to mouse brain expression level and genome baseline in prioritizing de novo variants in non-constraint genes with pLI<0.9, excluding known risk genes. pLI is excluded from the comparison because it is used in stratifying non-constraint genes. (C) Enrichment compared in top ranks by each metric. (D) Precision and true positives comparison.

### A-risk informs prior estimation in autism risk gene discovery

TADA and extTADA [39, 40] are empirical Bayesian methods used in previous genetic studies of autism[8, 20] to identify candidate risk genes based on burden of *de novo* variants. A key feature of such empirical Bayesian method is that it estimates parameters of priors, including mean relative risk (***R***) and prior probability (**π**) of being a risk gene, from the data. We reasoned that metrics associated with plausibility of autism risk, such as A-risk and gene constraint (pLI), could be used to improve prior estimation in an empirical Bayesian framework. To this end, we stratified a total of 18663 protein-coding genes by A-risk score 0.4 and pLI cutoff 0.9, resulting in 4 quadrants of genes (Supplementary Figure 5A): 1195 constrained genes with high A-risk score (quadrant A), 1842 constrained genes with low A-risk score (quadrant B), 1444 non-constrained genes with high A-risk score (quadrant C) and 14182 non-constrained genes with low A-risk score (quadrant D); then we estimated prior parameters by extTADA in each quadrant of genes, using previously reported *de novo* LGD and D-mis variant data from 8838 trios[6, 20–24]. Consistent with previous simulation[6], in unstratified analysis, **π** is about 0.04, corresponding to 750 risk genes in total. In stratified analysis, **π** decreases from quadrant A to quadrant D (Supplementary Figure 5B). Constrained genes stratified by A-risk ≥0.4 in quadrant A have greater **π** and ***R*** than genes with low A-risk scores in quadrant B (Supplementary Figure 5C). Genes in quadrant C and D have similar **π**, but quadrant C genes have a substantially greater ***R*** that D genes. Overall, A-risk informs the estimation of those priors in both constrained and non-constrained genes.

extTADA calculates a Bayes factor (BF) and posterior probability of association (PPA) for each gene, and then converts PPA to FDR (false discovery rate) to identify candidate risk genes. Common FDR procedures are designed to control the proportion of false positives among discoveries. However, with a large number of known risk genes ranked among the top by PPA, the estimated FDR of novel genes will be smaller than their true values, considering the true FDR of known genes is 0. This will lead to inflation of the support for novel candidate genes [41]. To address this issue, we excluded 90 known genes with SFARI gene score 1 or 2 in FDR estimation (Supplementary Table 6). The stratified analysis yielded 71 candidate genes passing FDR ≤0.1, whereas unstratified analysis yielded 44 genes. Among these genes, 38 were identified exclusively by the stratified approach, 11 were exclusively found by the unstratified approach, and 33 were shared (Figure 3). Previous studies have shown that autism risk genes are often pleiotropic and implicated in other neurodevelopmental disorders (NDD) [20, 42–44]. We obtained candidate NDD genes from a recent study[41] to seek support of the candidate autism genes. Among the 38 genes identified only in stratified approach, 13 are significantly implicated with NDD. In contrast, only 1 out of the 11 unstratified-exclusive genes is implicated with NDD (Supplementary Figure 6 and Supplementary table 7). Among the candidate genes that are also implicated with NDD, several are notable with additional support from other studies on autism or syndromes with autistic features, such as *NR2F2, NR4A2, HNRNPU, CLCN4,* and *PRKAR1B* **(Table 2)**. Candidate risk genes located in quadrant C, such as *GIGYF1* and *PRKAR1B,* are among the small number of candidate genes that are not constrained (pLI ~ 0).

**Figure 3.**
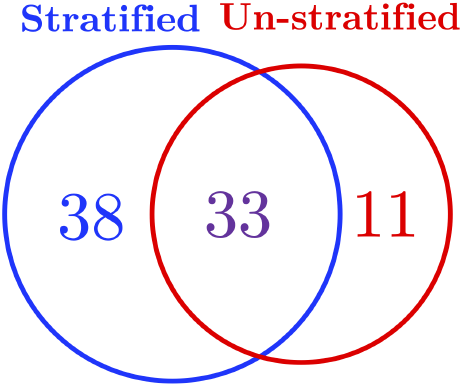
Stratified extTADA analysis by A-risk and pLI identifies more candidate risk genes of autism. The numbers in the Venn diagram show the number of genes identified by stratified analysis exclusively (38), by unstratified analysis exclusively (11), and by both approaches (33).

**Table 2.** Notable candidate risk genes by stratified extTADA analysis.

| Gene Symbol | pLI | A-risk | Gene quadrant | # of LoF, Dmis | FDR | NDD significant genes [41] | Additional support |
| --- | --- | --- | --- | --- | --- | --- | --- |
| *NR2F1* | 1 | 0.68 | A | 1, 1 | 0.07 | TRUE | Bosch-Boonstra-Schaaf Optic Atrophy Syndrome with autistic manifestation [45] |
| *NR4A2* | 1 | 0.43 | A | 1, 1 | 0.09 | TRUE | Levy 2018 [46] |
| *CLCN4* | 1 | 0.59 | A | 0, 3 | 0.015 | TRUE | Raynaud-Claes syndrome (OMIM 300114) with autistic features |
| *PRKAR1B* | 0.18 | 0.43 | C | 1, 2 | 0.06 | TRUE | Additional damaging variants in Ruzzo 2019 [47] |
| *GIGYF1* | 0 | 0.56 | C | 5, 0 | 1e-5 | TRUE |  |
| *HNRNPU* | 1 | 0.48 | A | 1, 1 | 0.09 | TRUE | Mosaic mutations [48] |

### Autism risk genes are highly expressed in deep-layer excitatory neurons in cortex

Previous studies have investigated autism risk by cortex laminar architecture. However, studies based on co-expression analysis [15, 16] or neurochemical experiments [49, 50] reported conflicting conclusions, that either deep or superficial layers of cortex are associated with autism. These early studies were based on a small number of high-confidence autism risk genes. Here we revisit the question with a much larger list of high-confidence candidate genes and single cell RNA-seq data. We obtained a single-nucleus RNA-seq data set of the middle temporal gyrus (MTG) of adult human cortex with clear laminar layer information [51]. The expression level of those 90 SFARI score 1 or 2 genes and 71 novel candidate risk genes is shown in the heatmap in Figure 4A. Hierarchical clustering based on the expression data forms four major clusters of genes. Genes in cluster 1 show very little expression in most cell types, except that *TBR1, RORB, MEIS2, PTCHD1, FEZF2* and *NR4A2* are sparsely expressed in subtypes excitatory neurons and *RELN* and *PCDH19* are highly expressed in subtypes of inhibitory neurons. Cluster 2 genes have more specific expression in deep-layer excitatory neurons. Genes in cluster 3 are expressed more widely in neuronal cell types with even higher expression in excitatory neurons at deep layers of MTG. Genes in cluster 4 have high expression in almost all the neuronal cell types in MTG. Mapping quadrant gene groups defined by A-risk and pLI into those 4 distinct expression clusters reveals that both cluster 3 and 4 are dominated by quadrant A genes (33 out of 47 genes and 29 out of 32 genes, respectively). Cluster 2 contains the largest portion of quadrant C genes (10 out of 16 genes, Figure 4B). Consistent with pLI value distribution, a larger fraction of genes in cluster 2 have higher observed to expected (O/E) ratio of LoF mutations in gnomAD (genome aggregation database)[52] compared to genes in other clusters (Figure 4C). Overall, excitatory neurons project from or to deep layers have high expression of the largest subset of known and candidate risk genes.

**Figure 4.**
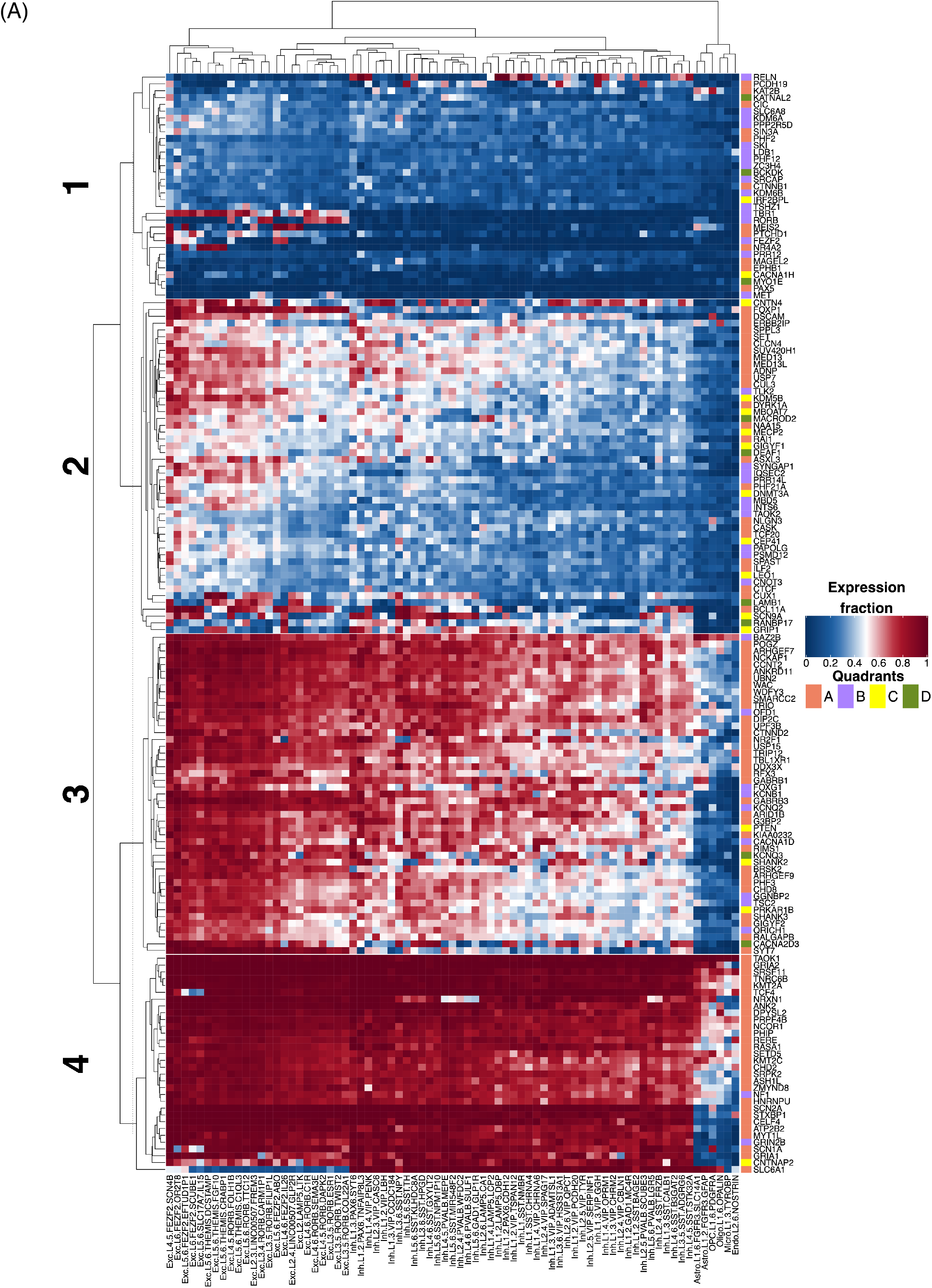
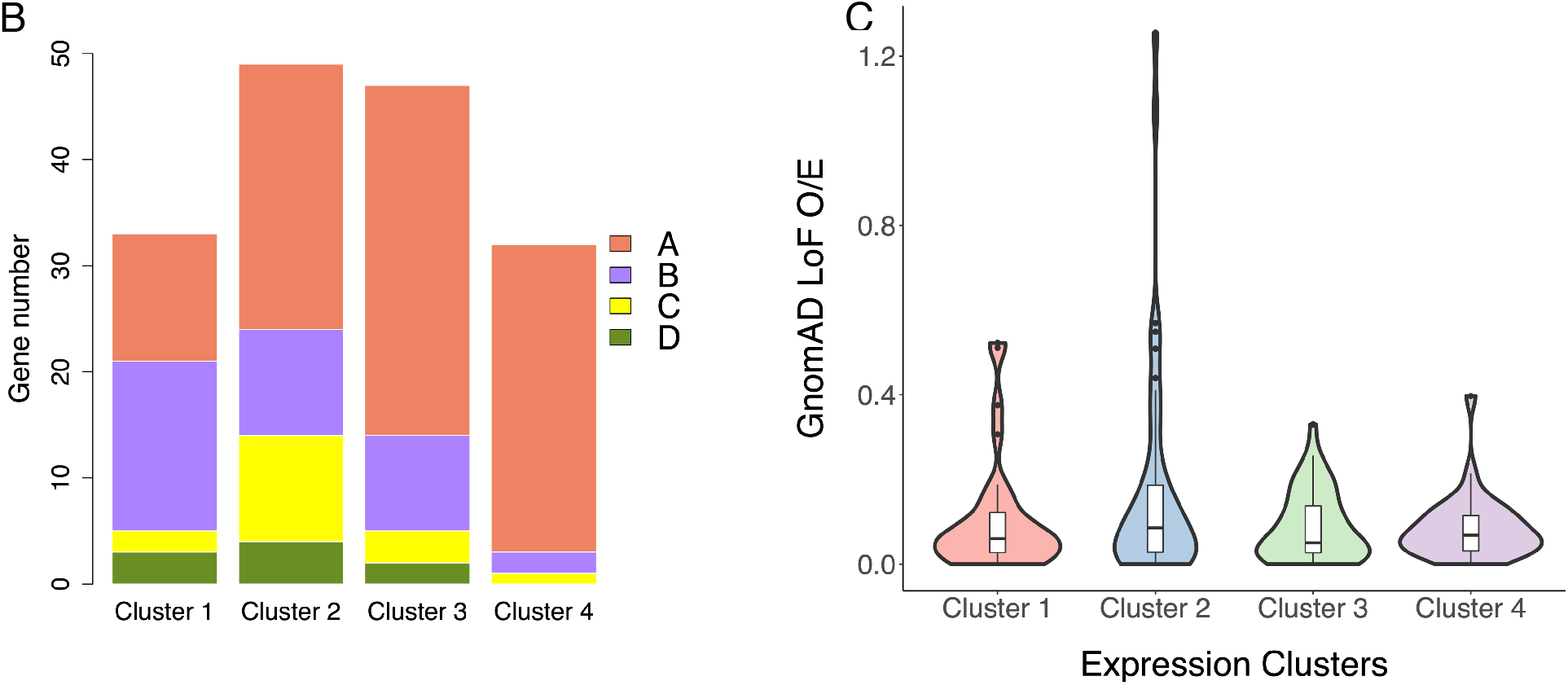
Most autism risk genes have high expression in deep-layer excitatory neurons in prefrontal cortex. (A) Hierarchical clustering 90 known autism risk genes and 71 novel candidate genes by expression level in cell types from adult cortex middle temporal gyrus (MTG) with laminar information. Genes (shown in rows) form 4 major clusters, labeled from 1 to 4 on the left. The dash line marks the height cutting the hierarchical tree. Cell types are clustered as well and are labels in the format as “major cell type.located layers.marker genes”. Exc, excitatory neurons. Inh, inhibitory neurons. Astro, astrocytes. OPC, oligodendrocyte precursor cells. Oligo, oligodendrocytes. Micro, microglia. Endo, endothelial cells. The color (blue to red) of the heatmap indicates expression level of a gene in the cell type, calculated as the fraction of cells that have ≥1 UMI mapped to the gene in the cell type. Almost all genes in cluster 1 have low expression in all cell types. Most genes in cluster 2 are specifically expressed in excitatory neurons in deep layers (layer 4 to 6). Cluster 3 genes are highly expressed in deep excitatory neurons and have expression in most of neuronal cell types. Cluster 4 genes are highly expressed in almost all neuronal cell types. Quadrant gene groups stratified by Frisk and pLI are labeled by the color bar on the right side with A, B, C and D represented by orange, purple, yellow and green. (B) Number of known or candidate risk genes from quadrant gene groups in each expression clusters. Cluster 1 is enriched with quadrant B genes (high pLI and low A-risk); cluster 2 is enriched with quadrant C genes (low pLI and high A-risk); cluster 3 and 4 are enriched with quadrant A genes (high pLI and high A-risk). (C) The distribution of observed over expected (O/E) number of loss of function variants in gnomAD database in the 4 expression clusters. Cluster 2 genes have a broad distribution of O/E. Genes in other clusters have generally small O/E.

The heatmap of expression fraction in the same order of genes using the two fetal data sets in our model are shown in Supplementary Figure 7. There is no layer information with the fetal data. Nevertheless, the expression patterns of candidate risk genes in the two fetal data sets generally follows the organization in the adult cortex data, especially for fetal prefrontal cortex. Additionally, 14 out of 24 cluster 1 genes with little expression in adult cortex neuronal cells have fraction expression ≥0.5 in at least one cell type in fetal prefrontal cortex, suggesting a dynamic temporal specific expression of those candidate risk genes.

## Discussion

In this study, we developed a new method, “A-risk”, to predict plausibility of autism risk genes based on single-cell expression patterns in human fetal midbrain and prefrontal cortex. A-risk was trained using known autism genes. A-risk score reflects the similarity of the cell-type-specific expression pattern of a gene to known autism genes in aggregation. It achieves superior performance in prioritizing *de novo* risk variants, especially in genes that are less intolerant of loss of function variants. Furthermore, A-risk is complementary with gene constraint metric (pLI) for improving estimation of priors using an empirical Bayesian association method. Applying it to published *de novo* variant data, we identified 71 novel candidate risk genes, an increase of 27 genes over the results using the same statistical method without stratification of genes by either A-risk or pLI.

Both inhibitory and excitatory neurons in the prefrontal cortex strongly contribute to A-risk prediction during fetal stages, consistent with previous theory of excitatory and inhibitory imbalance in the prefrontal cortex disrupting neural communication[29, 53]. GABAergic inhibitory neurons in midbrain have been identified as the most significant contributing feature to A-risk prediction, implicating a potential role of midbrain in autism pathogenesis that has been understudied.

Early functional and co-expression network studies [11, 15] based on a small number of high-confidence autism risk genes have revealed convergence on excitatory neurons in deep-cortical layers, however, another co-expression network analysis [16] found significance in excitatory neurons in superficial cortical layers. With a much larger number of high-confidence risk genes, we revisited the role of neuronal cell types in six different cortical layers. Based on a large single nuclei RNA-seq data set from adult cortex, we observed that deep-layer excitatory neurons have high expression of the vast majority of known and candidate autism risk genes, while other neuronal types or neurons in superficial layers have high expression of a much smaller subset of these genes. Since the excitatory neurons residing in layer 5 or 6 of cortex extend their axons into other regions of brain and communicate between cortex and other critical regions [29, 54], disruption of deep-layer excitatory neurons more likely affects signal transmission and communication across different brain regions. Taking account of gene mutation intolerance (pLI) and expression similarity to known autism genes (A-risk), the candidate risk genes with high A-risk but low pLI (i.e. quadrant C), such as *GIGYF1* and *MBOAT7,* are much more likely to have specific expression in deep-layer excitatory neurons. Interestingly, a recent study [20] showed *GIGYF1* was the most autism-specific gene among all candidate autism risk genes based on frequency of disruptive *de novo* variants in either autistic or severe NDD cohorts. This suggests an association of deep-layer excitatory neurons and autistic conditions that do not involve severe NDD conditions such as intellectual disabilities. We expect that this hypothesis will be tested in future studies with independent high-resolution single cells or neural circuit expression data, larger set of high-confidence risk genes, and autism cohorts with comprehensive NDD phenotyping.

The majority of genes in quadrant C are located in expression cluster 2, where a higher proportion of genes shows increased observed to expected (O/E) ratio of LoF mutations, suggesting quadrant C genes are less intolerant to LoF mutations or may be incompletely penetrant. The genes that have high A-risk and high pLI (quadrant A) are more likely to have high expression in a wide range of cell types. Candidate risk genes in cluster 1, among which 16 genes out of 33 in total have high pLI but low A-risk (quadrant B), have sparse expression in adult cortex but more expression in fetal prefrontal cortex, indicating those autism risk genes can take effect at limited time points and places.

A-risk directly utilized single-cell transcriptomic data as the input of the machine learning model to learn expression patterns from known risk genes. Expression patterns inferred from single-cell RNA-seq data have better resolution than bulk sequencing data with fine-grained cell-type heterogeneity and developmental temporal information. To integrate transcriptomic information in risk gene discovery in a principled way, we used A-risk in an empirical Bayesian framework to improve prior estimation based on genetic data. This approach yielded 27 more candidate risk genes than the original Bayesian approach using only genetic data. With increased sample sizes in the future, A-risk can also be used as informative covariates to improve FDR estimation [55, 56] in frequentist approaches for risk gene discovery.

A-risk is currently limited by the availability of comprehensive single-cell expression profiles across all critical human brain regions and developmental stages. Profiling neuron cells is uniquely challenging since the information in extended projections and axons can be lost during sample preparation in single-cell RNA-seq. Even though the data we used in the A-risk model is from fetal stages, when extended axons of neurons have not been prolonged, we should still interpret with consideration that there could be some genes missed in the data. New single-nucleus RNA-seq and subcellular transcriptomic profiling techniques and data sets from ongoing projects such as Allen Brain Institutes [57] and Human Cell Atlas [58] will help to address this issue [59, 60]. Additionally, A-risk is a supervised learning approach, and inevitably it biases towards genes with similar expression patterns to known risk genes in the training. Unsupervised approaches could assist in addressing the problem. Finally, abundant and specific expression is not sufficient to define a gene as a risk gene. Other factors such as functional redundancy [61] and protein complex formation [62] that determine whether a high-expression gene is a bottleneck in a system, also play a role in the genetic impact. Future studies can consider those factors with single-cell expression profiles to improve accuracy of prediction.

## Methods

### Data collection and preprocessing

In this study, we integrated human fetal brain single-cell RNA-seq data from two publications: (1) midbrains from 6 to 11 weeks [26] and (2) prefrontal cortexes from gestational weeks 8 to 26[27]. To integrate these two data sets, first, we obtained the UMI counts of single cells from their published data. Second, we directly utilized the cell type clusters and time points documented in the publications and calculated the expression fraction of each gene in each cell type at a particular time point. We combined each individual cell type and time point together to generalize one feature in the integrated data. The expression fraction is defined as, for a particular gene in a cell type at a developmental time point, the number of cells having the gene expressed (UMI >= 1) divided by the total number of cells grouped in the cell type. *La Manno et al., 2016[26]* reported 26 cell types across 6 developmental time points, including an unknown cell type (“Unk”) where those cells cannot be assigned to any known clusters. We excluded Unk cells in the analysis. *Zhong et al., 2018* [27] reported clustered 35 cell types through 9 time points. Furthermore, we also excluded cell types with fewer than or equal to 10 cells. In total, we compiled 95 features in the combined data set, including 47 from *La Manno et al., 2016* [26] and 48 features from *Zhong et al., 2018* [27].

We obtained known autism risk genes with score of 1 or 2 in the SFARI database[7] (https://gene.sfari.org/database/human-gene/, version released on 08/29/2019) as the positives for model training. For the negatives for model training, we collected genes harboring at least 1 *de novo* LGD variant in controls from an exome-sequencing study on autism[6]. Two genes *(KDM5B* and *CACNA1H)* are present in both the initial positive and negative sets. We removed these 2 genes from the negative set. In total, we compiled 88 genes in the positive training data set and 154 genes in the negative training data set. The full list of training genes is available in Supplementary table 1.

### Machine learning approaches to predict autism risk genes

We trained a supervised machine-learning method, gradient boosting tree, using the training gene set and features derived from single cell data sets. To implement the gradient boosting tree machine, we used the python package “sklearn.ensemble.GradientBoostingClassifier” with parameters of “n_estimators” as 300, “learning_rate” as 0.05 and “max_depth” as 1. We assessed the performance of the model by 5-fold cross validation. In each cross validation, the model randomly selected 20% of the training gene set to serve as a test set for validation and the rest of the genes were used to train the model. We implemented the python package “sklearn.metrics.roc_curve” to calculate the true positive rate, false positive rate, and to plot the ROC curve and calculate AUC values. After training, we predicted the probability for each protein-coding gene in the genome being a positive gene (i.e. plausibility for being an autism risk gene) by the trained model. The final A-risk score is the average probability from the 5-fold training and prediction. The complete A-risk score is available in Supplementary table 2.

“Feature importance” is derived from the gradient boosting tree model using the function “feature_importances_”. The final feature importance value for each selected feature is the average from the 5-fold training and prediction. All selected features with non-zero feature importance are listed in Supplementary table 2.

### Comparison of A-risk to other metrices in prioritizing *de novo* LGD variants

We tool two approaches to compare the ability of A-risk and other metrics in prioritization of *de novo* variants. With each metric, we first rank all genes; then in all genes above a certain rank threshold (e.g. 1000, 1500, 2000, etc), we estimated the “enrichment of *de novo* variants”, “precision”, and “true positives”. The formulae to compute these estimates are as following:

For any gene ***i***, the number of expected *de novo* variants in each gene, ***E_i_***, was calculated as:

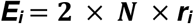

where ***N*** is the number of trios in the compiled data sets and ***r_i_*** is gene-specific background mutation rate. Here we tested on *de novo* gene-likely disrupting (LGD) variants and deleterious missense (D-mis) variants (Figure 2). LGD variants include nonsense, frameshift and canonical splice site mutations and D-mis variants are defined as variants with REVEL (the Rare Exome Variant Ensemble Learner) score >= 0.5[37]. For each gene, ***r_i_*** is the sum of background mutation rate of LGD mutations plus D-mis mutations.

The background mutation rate per gene of each mutation type was obtained from a previous described mutation model [35, 36]. Briefly, the seven-nucleotide sequence context was used to determine the probability of each base in mutating to each other possible base. Then, the mutation rate of each functional class in each gene was calculated by adding up point mutation rates in the longest transcript. The rate of frameshift indels was presumed to be 1.25 times the nonsense mutation rate and the rate of genes located on chromosome X is further adjusted according to female-to-male ratio in the *de novo* data set [63].

For a set of genes, the enrichment of *de novo* variants, ***D,*** was calculated as:

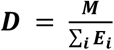

where ***M*** is the total number of observed *de novo* LGD or D-mis variants in this gene set. In this study, we compiled results from multiple whole exome studies on autism spectrum disorders, including total of 8838 trios from Simons Simplex Collection (SSC) [6], Autism Sequencing Consortium (ASC) [20], SPARK Pilot [21], MSSNG [22], Takata et al., 2018 [23] and Chen et al., 2017 [24] cohorts.

For any gene set, the number of detected true positives, ***TP***, was calculated as:

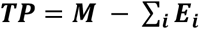

For any gene set, the precision (positive predictive value), ***PPV***, was calculated as:

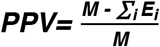

For each metric (A-risk, pLI etc.), a set of genes were selected based on the rank of genes by each individual metric, such as top 1000 genes or top 2000 genes, etc. The genome baseline is defined by all the genes in the genome. For the first estimate, enrichment of *de novo* variants, ***D***, was calculated for any set of top-ranked genes, and then enrichment values were plotted and compared, as shown in Figure 2A. For the second estimate, the number of detected true positives, ***TP***, and the precision (true discovery rate), ***PPV***, were calculated for any set of top-ranked genes. ***TP*** and ***PPV*** were plotted and compared, as shown in Figure 2B. Recall would be calculated as *R* = ***TP/N***, where *N* is the total number of true positives (*N*). Since *N* is unknown but a constant, ***TP*** is proportional to *R*. Therefore, we use *TP* as a proxy of recall. To avoid inflation of A-risk performance, we excluded all the known autism risk genes used in A-risk training during calculation of all above estimates. Although there are different numbers of genes predicted by each method, we compared all the methods with 18663 protein-coding genes, replacing missing scores with the median of each corresponding metrices.

To exam the potential of A-risk in prioritizing *de novo* variants in non-constrained genes, we limit the estimates on genes with pLI score <= 0.9 in each top rank of genes (Figure 2C and 2D). We excluded pLI as a metric for comparison in those figures since pLI was used to stratify constraint and non-constraint genes. Furthermore, we also compare A-risk with the other two metrices D-score and Krishnan 2016 (Supplementary Figure 4).

### Application of A-risk in stratified risk-gene discovery analysis

In this analysis, we used an empirical Bayesian model of rare-variant genetic architecture, extTADA (Extended Transmission and *de novo* Association)[39, 40], which can estimate mean effect sizes and risk-gene proportions from the genetic data to identify autism candidate risk genes. To inform the prior estimation, we stratify all protein-coding genes into 4 quadrants by A-risk score 0.4 and pLI 0.9. Specifically, quadrant A consists of genes with A-risk >=0.4 and pLI >=0.9. Genes in quadrant B are in A-risk <0.4 but pLI >=0.9. Genes in quadrant C have A-risk >=0.4 but pLI <0.9, and the rest of the genes are assigned to quadrant D. We applied the extTADA model to each quadrant of genes to estimate the priors and calculate posterior probability of association (PPA). Then we combined the posterior probability of the 4 quadrants together to calculate a final FDR (false discovery rate). To make FDR estimation of novel risk genes more accurate, we excluded known autism risk genes used in training A-risk model in FDR calculation, as most of these genes are ranked in top by PPA and including them in FDR calculation with deflate FDR values of novel risk genes. In parallel, we did the same analysis in all genes without stratification by A-risk or pLI. We used the same *de novo* variant data from 8838 trios and background mutation rate data as described above. Final results summarizing both stratified and unstratified analysis are available in Supplementary Table 3.

### Expression pattern clustering of known and candidate autism risk genes

We compiled the 71 novel candidate risk genes that pass FDR <=0.1 in stratified extTADA analysis together with 90 known risk genes and investigated the expression pattern of all those risk genes in a single-cell RNA-seq data of adult human cortex[51]. The data was preprocessed as described above and the expression fraction for each cell type was pre-computed from read-count data downloaded from the publication. Hierarchical clustering was performed using “ComplexHeatmap” package in R based on “Euclidean distance” and the heatmap (Figure 3) was drawn by the “heatmap” function built in the package.

### Data and code availability

Single-cell RNA-seq data in *La Manno et al., 2016* and *Zhong et al., 2018* are downloaded from Gene Expression Omnibus (GEO) with accession number as GSE76381and GSE104276 respectively. Data reported from *Hodge et al., 2019* is downloaded from https://portal.brain-map.org/atlases-and-data/rnaseq.

A-risk model training and prediction is implemented using python package “sklearn.ensemble.GradientBoostingClassifier”: https://scikit-learn.org/stable/modules/generated/sklearn.ensemble.GradientBoostingClassifier.html.

The python script and processed data for A-risk model is available on GitHub: https://github.com/ShenLab/A-risk.

## Acknowledgements

This work was supported by grants from the National Institute of General Medical Sciences (R01 GM120609), the National Heart, Lung and Blood Institute (R03 HL147197) and Simons Foundation Autism Research Initiative (606450). We thank Drs. Wendy Chung, Nicholas Tatonetti, and Chaolin Zhang for helpful discussions.

