## Supplementary Figures and Tables for "Dissecting Autism Genetic Risk Using Single-cell RNA-seq Data"

Chen S.  et al

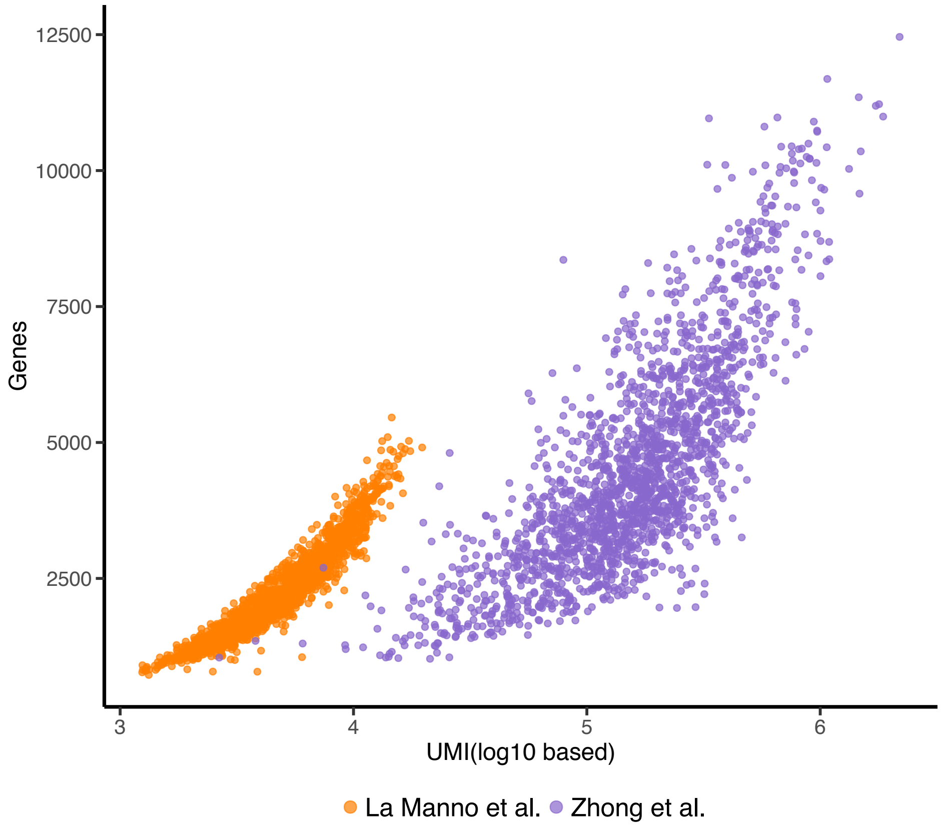


**Supplementary Figure 1.** Quality of single cell RNA-seq data. The number of log10 based UMIs in each cell from the two data sets against the number of genes detected. The detected genes are defined as genes with larger or equal to 1 UMI. The midbrain data[1] has more genes detected than the prefrontal cortex data[2] given the same number of UMIs.

(A)

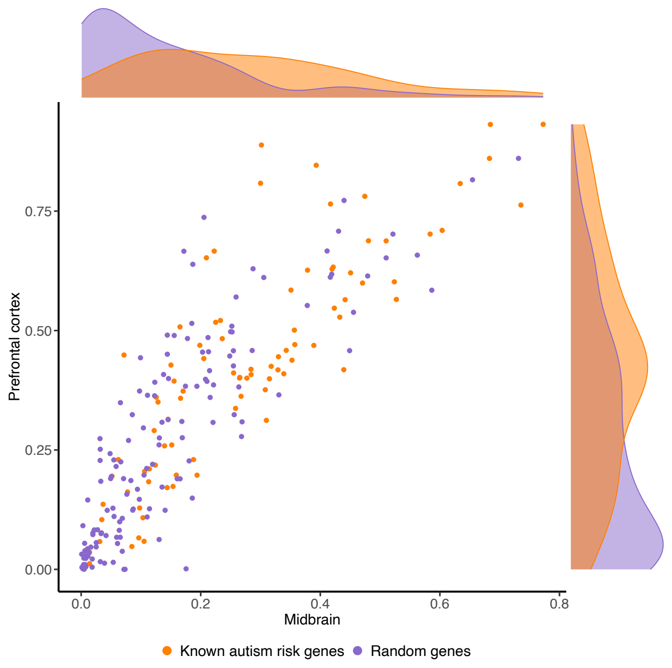


(B)

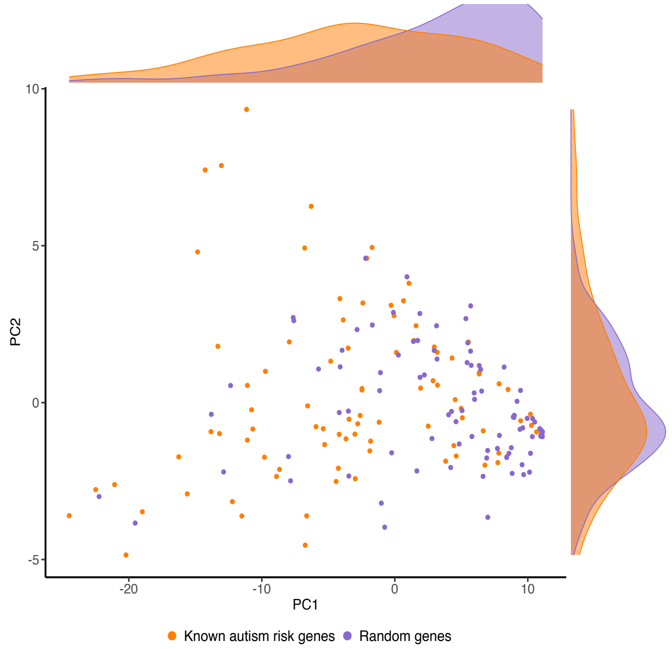


**Supplementary Figure 2.** Different expression pattern of known autism risk genes and random genes in fetal midbrain and prefrontal cortex. A. The expression distribution of known autism risk genes and random genes in fetal midbrain and prefrontal cortex. B. PCA analysis of fraction expression of known autism risk genes and random genes. The density plots along axes shows the difference of known risk genes and random genes in expression level or PCA scores.

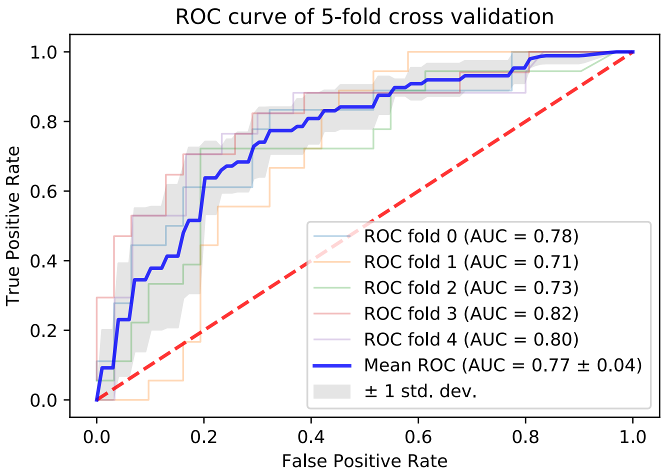


**Supplementary Figure 3.** Training of A-risk: performance in cross-validation and importance of cell types and time points to the model. A. ROC curves of 5-fold cross validation using training data, where the training samples are divided as 80% for training and 20% for validation. The blue curve is the average of the 5 curves and the grey band in the background marks the interval between the left and right first standard deviation.

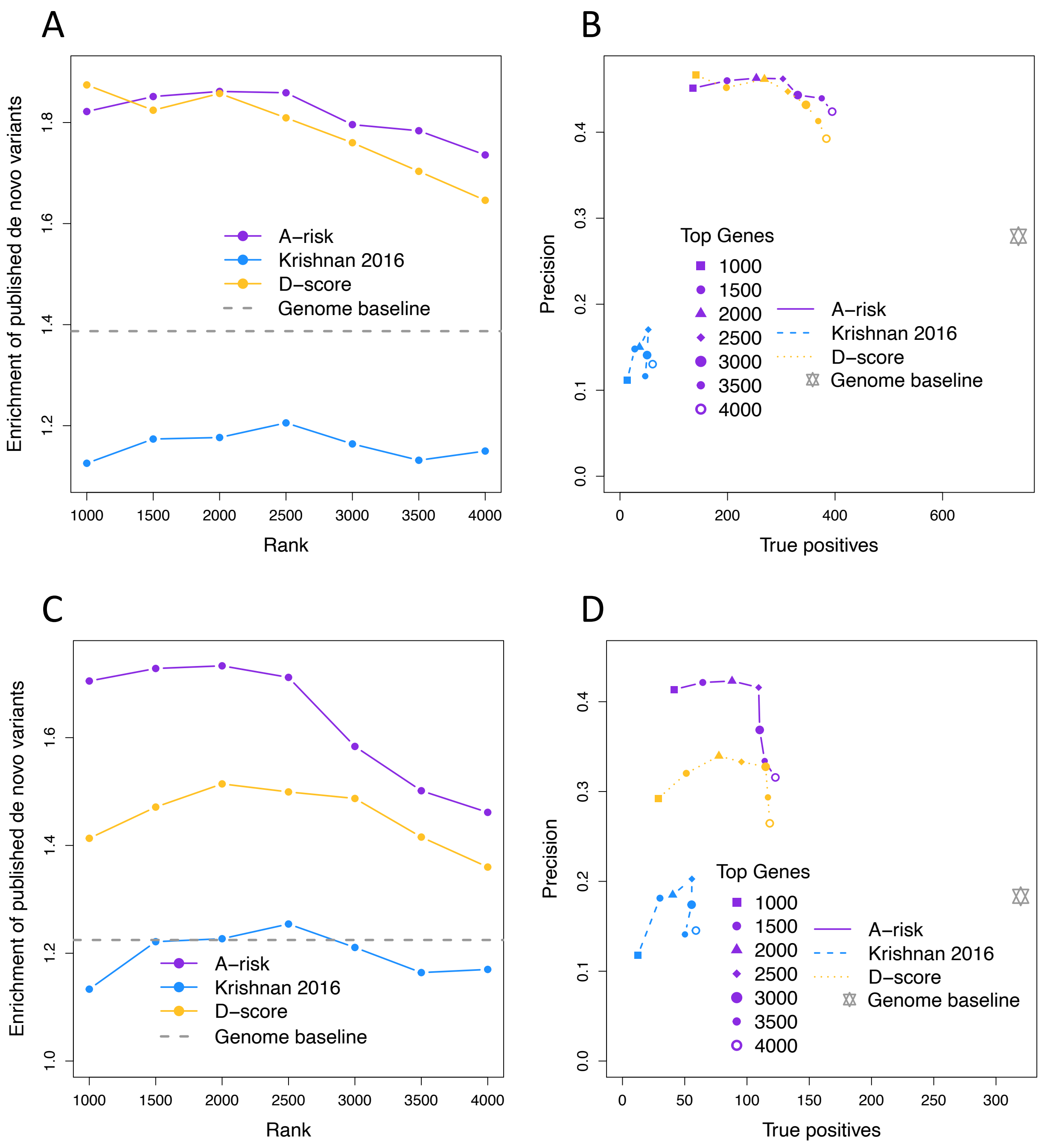


**Supplementary Figure 4.** A-risk has better performance than other two methods in prioritizing de novo variants. A-B, Compare A-risk to Krishnan 2016[3] and D-score[4] in enrichment, precision and true positives of de novo LGD and D-mis variants prioritized in top ranks by each method, excluding all known risk genes. C-D, Compare the three methods in non-constraint genes stratified by pLI < 0.9, excluding all known genes.

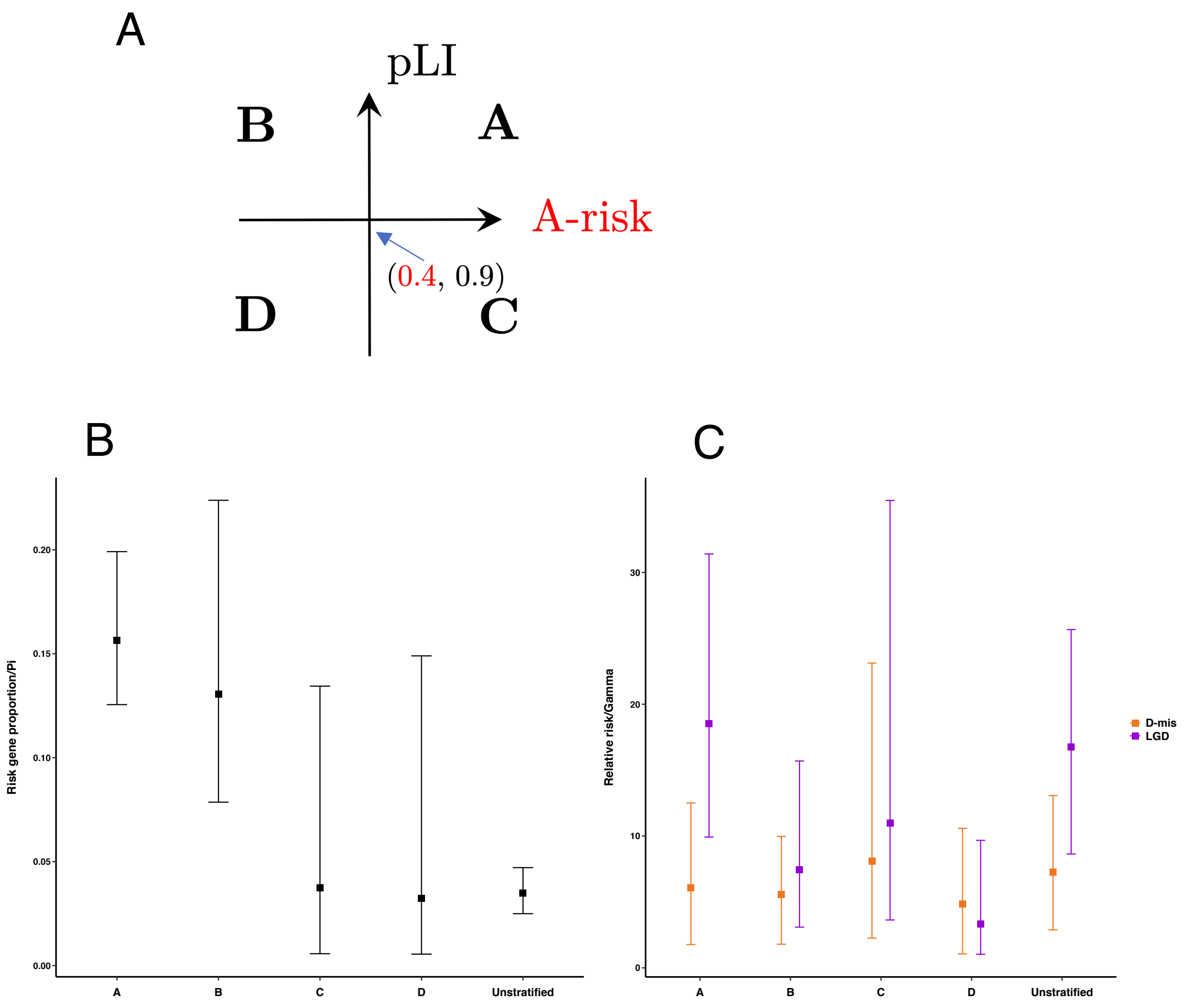


**Supplementary Figure 5.** Prior estimation in stratified extTADA analysis. Panel (A). gene groups defined by pLI and A-risk: A: pLI≥0.9 and A-risk≥ 0.4; B: pLI≥0.9 and A-risk <0.4; C: pLI<0.9 and A-risk≥0.4; D: pLI<0.9 and A-risk<0.4.
Panel (B). Risk gene proportions (π) in stratified gene groups estimated from MCMC. Modes are indicated by small boxes in the middle and the upper and lower bars indicate 95% confidence intervals. Panel (C). Relative risks (γ) of genes in each stratified group estimated from MCMC. Relative risks estimated separately from LGD and D-mis variant data, labeled by purple and orange respectively.

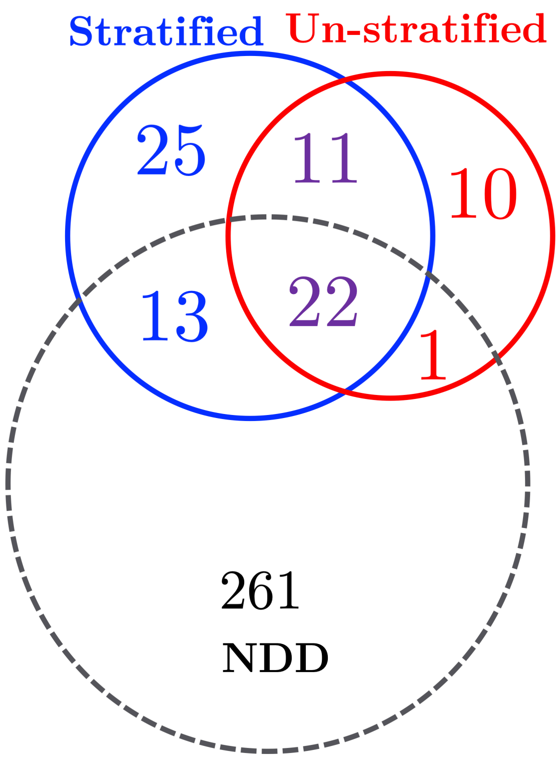


**Supplementary Figure 6.** Additional support of candidate novel autism risk genes identified by stratified or unstratified extTADA analysis with significant genes in neurodevelopmental disorders (NDD) identified by Kaplanis et al 2020[5]. Among 33 genes identified by both stratified and unstratified extTADA, 22 (67%) are implicated with NDD; 13 genes out of 38 (34%) identified exclusively by stratified extTADA are implicated with NDD, whereas only 1 out of 11 (9%) exclusively identified by unstratified extTADA is associated with NDD.

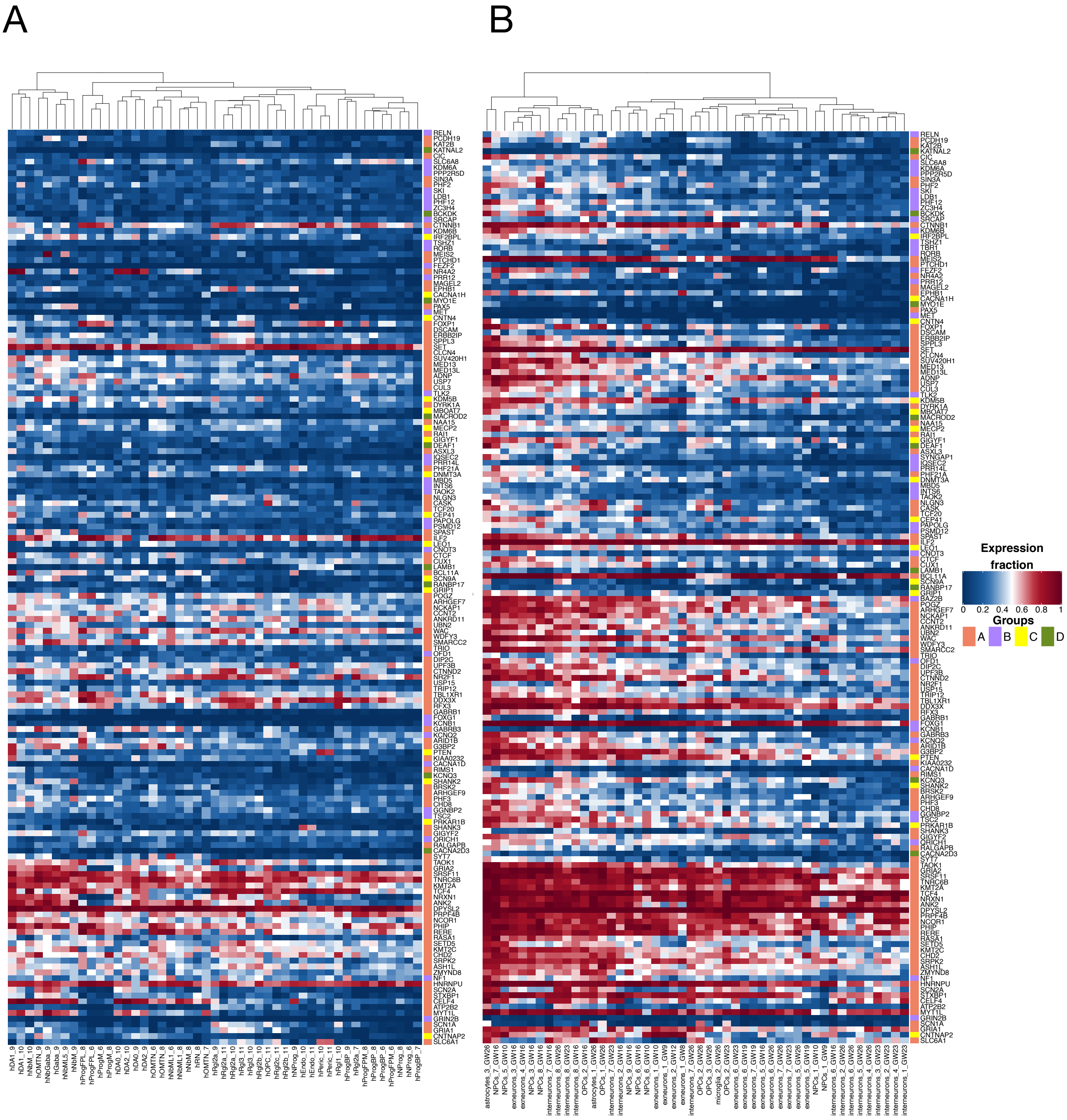


**Supplementary Figure 7.** Heatmap of expression level of known and candidate risk genes in fetal midbrain[1] (A) and prefrontal cortex[2] (B). Row orders are arranged as same as Figure 3. Cell types in midbrain are labeled as "h(human)cell type names_week" and cell types in prefrontal cortex are labeled as "major cell type name_sub clusters_gestational weeks", in concordance with original data. DA, dopaminergic neurons. NbM, medial neuroblast. OMTN, oculomotor and trochlear nucleus. NbGaba, neuroblast GABAergic. Gaba, GABAergic neurons. NbML, mediolateral neuroblasts. ProgFPL, progenitor lateral floorplate. ProgM, progenitor midline. RN, red nucleus. Rgl, radial glia-like cells. OPC, oligodendrocyte precursor cells. NProg, neuronal progenitor. Endo, endothelial cells. Peric, pericytes. ProgBP, progenitor basal plate. ProgFPM, progenitor medial floorplate. NPCs, neural progenitor cells. Exneurons, excitatory neurons.

---

**Supplementary Table 5.** Summary of publication sources of *de novo* variants data.

| Cohort label | Number of unique cases | Publication |
|---|---|---|
| ASC | 3625 | Satterstrom et al., 2019[6] |
| De Rubies | 421 | De Rubeis et al., 2014[7] |
| SSC | 2501 | Iossifov et al., 2014[8] |
| SPARK pilot | 465 | Feliciano et al., 2019[9] |
| MSSNG | 1529 | Yuen et al., 2017[10] |
| JPASD | 232 | Takata et al.,2018[11] |
| ACE | 65 | Chen et al.,2017[12] |
| Total | 8838 | |

The following tables are included in a separate spreadsheet:

**Supplementary Table 1.** Cell types in which the expression level of genes was used as features for model training.
**Supplementary Table 2**. Known risk genes and random genes used in model training.

**Supplementary Table 3**. A-risk score of all protein-coding genes predicted by the model.

**Supplementary Table 4.** Feature important estimated by the gradient boosting method.

**Supplementary Table 6.** extTADA results of all protein-coding genes.
**Supplementary Table 7.** Risk genes detected by stratified and unstratified extTADA.
